# *In vitro* analysis of *Fusobacterium Polymorphum* from oral leukoplakia patients identifies “high-risk” isolates

**DOI:** 10.1101/2025.10.01.679755

**Authors:** Doaa E. El-Hadedy, Gary Moran, Nezar N. Al-Hebshi

## Abstract

**Introduction:** Mounting evidence links *F. nucleatum*, particularly subsp. *polymorphum* (now classified as *F. polymorphum*) to oral squamous cell carcinoma (OSCC). However, not all studies support this association, and its ubiquitous presence in health complicates its role as a driver of disease, raising a fundamental question: Can some isolates be pathogenic “high risk” while others are not? Here, we investigated the carcinogenic potential of clinical isolates of *F. polymorphum* from oral dysplastic lesions.

**Methods:** Sixteen, fully sequenced *F. polymorphum* isolates from healthy subjects (6 isolates) or patients with mild epithelial dysplasia (3 isolates), moderate dysplasia (3 isolates) or severe dysplasia/OSCC (4 isolates) were included. The isolates were assessed for their effect on the proliferation, migration, invasion, transcriptome and cytokinome of dysplastic oral keratinocytes (DOK). The isolates were also subject to RNA-seq and secreted amyloid FadA analyses.

**Results:** Although genetically indistinguishable, the isolates differed markedly in their effects on dysplastic oral keratinocytes (DOK) with isolates from dysplastic lesions demonstrating enhanced proliferation, migration, and invasion of DOK in proportion to the dysplasia severity of their clinical origin. Strikingly, amyloid-like FadA levels were also significantly higher in dysplasia-associated “high-risk” isolates and correlated with their pro-carcinogenic effects. RNA-seq further showed upregulation of heme acquisition genes in high risk isolates. Transcriptome and cytokine profiling of DOK revealed a uniform pro-inflammatory response across all isolates, independent of origin, but genes and pathways related to proliferation correlated with dysplasia severity. Supporting this, 45% of the most severity-correlated host genes identified *in vitro* were also differentially expressed in the same direction in tumors versus normal tissues in the TCGA OSCC dataset.

**Conclusion:** These results show, for the first time, that clinical oral isolates of *F. polymorphum* vary in their carcinogenic properties, establishing the novel concept of “high-risk” versus “low-risk” isolates in oral carcinogenesis, potentially driven by genome-independent regulation.

## Introduction

Over the past decade, *F. nucleatum* has attracted significant attention for its potential as an oncobacterium in a growing range of cancers—particularly colorectal cancer (CRC), but also oral, esophageal and breast cancers (1-5). Multiple studies have demonstrated that higher levels of *F. nucleatum* in CRC tissues are correlated with advanced tumor staging, greater chemo-resistance, and worse patient outcomes (6-8). The bacterium is thought to contribute to a proinflammatory microenvironment, promoting cell proliferation, migration, and metastasis within tumors (9-13).

Mechanistically, the pathogenicity of *F. nucleatum* is intimately linked to its adhesive surface proteins, particularly those belonging to the type V secretion system such as Fap2 and FadA (12, 14). Fap2 binds to Gal-GalNAc residues (14), which are upregulated in CRC and breast cancer tissues, resulting in the induction of chemokines critical for inducing host cell migration such as IL-8 and CXCL1 (9). FadA, meanwhile, binds to E-cadherin on host cancer cells, activating β-catenin signaling pathways via Annexin A1, promoting carcinogenesis (12, 15). FadA forms amyloid-like structures that have been identified in diseased tissues in humans and to promote CRC progression in murine models (16).

The role of *F. nucleatum* in the initiation and progression of oral squamous cell carcinoma (OSCC) is less well studied. Studies frequently report increased abundance of *F. nucleatum* in OSCC tissues compared to healthy oral sites (1, 17, 18), with one study showing the proportion of *Fusobacterium* to rise with tumor progression, from roughly 3% in healthy controls to nearly 8% in advanced OSCC cases (18). *F. nucleatum* has also been found enriched in oral potentially malignant disorders (OPMD) such as leukoplakia, increasing in abundance with the degree of epithelial dysplasia (19, 20).

Meta-transcriptomic analyses have revealed that at tumor sites, *F. nucleatum* is metabolically hyperactive, with increased expression of pathways involved in iron acquisition, oxidative stress response, and proteolysis (21). Spatial studies using RNA sequencing and fluorescence in situ hybridization (FISH) have shown that *F. nucleatum* localizes to specific micro-niches within OSCC tissues, where it is associated with localized immunosuppressive effects (22). These areas show increased expression of ARG1 (arginase 1) and programmed cell death protein 1 (PD-1), both of which play pivotal roles in carcinogenesis (23, 24).

Notably, *F. nucleatum* is a ubiquitous and abundant oral bacterium even in health (25, 26), raising a fundamental question: what distinguishes its role in health from its involvement in carcinogenesis? Indeed, several studies failed to show an association between *F. nucleatum* abundance and OSCC or OPMD (27-30), suggesting that relative abundance alone is insufficient as a biomarker of disease. A more plausible explanation is that *F. nucleatum* isolates vary in their carcinogenic potential. We recently showed that the subspecies most commonly recovered from both normal and dysplastic oral mucosa is *F. nucleatum* subsp. *polymorphum* (31)—now classified as *F. polymorphum*. Although comparative genomic analysis revealed no distinct phylogenetic separation between isolates from health and disease, we observed considerable inter-individual variability in adhesin-encoding genes such as *fadA* and *fap2* (31). In subsequent functional assays, these isolates were found to induce migration and invasion of oral keratinocytes in an isolate-specific manner (32).

These findings underscore the need to move beyond community-level associations toward a deeper understanding of the functional diversity and pathogenic potential of clinical *F. polymorphum* isolates in oral carcinogenesis. Accordingly, the objective of this study was to investigate whether the carcinogenic potential of *F. polymorphum* isolates correlates with the severity of epithelial dysplasia of the lesions from which they are isolated.

## Methods

### *F. polymorphum* clinical isolate selection

Clinical isolates of *F. polymorphum* used here were recovered from healthy controls and patients with oral leukoplakia with varying levels of dysplasia attending the Dublin Dental University Hospital, as described by Crowley et al (31). The degree of epithelial dysplasia (mild, moderate or severe) was determined following biopsy by an experienced oral pathologist. The 16 isolates were selected from the collection to represent a spectrum of clinical origins: healthy patients (n=6), patients with mild epithelial dysplasia (n=3), moderate dysplasia (n=3) or severe dysplasia/OSCC (n=4) (**Fig 1A**). All *F. polymorphum* isolates were previously characterized by whole genome sequencing (31). The core and accessory genomes were identified with Panaroo (33) using the default settings. The core genome alignment file from this was used to produce a phylogenetic tree using Fastree (version 2.1.10) (34). Further gene annotation of the pangenome was carried out using EggNog (35). Genome-wide association analysis was carried out using SCOARY (version 1.6.16) (36).

**Figure 1.**
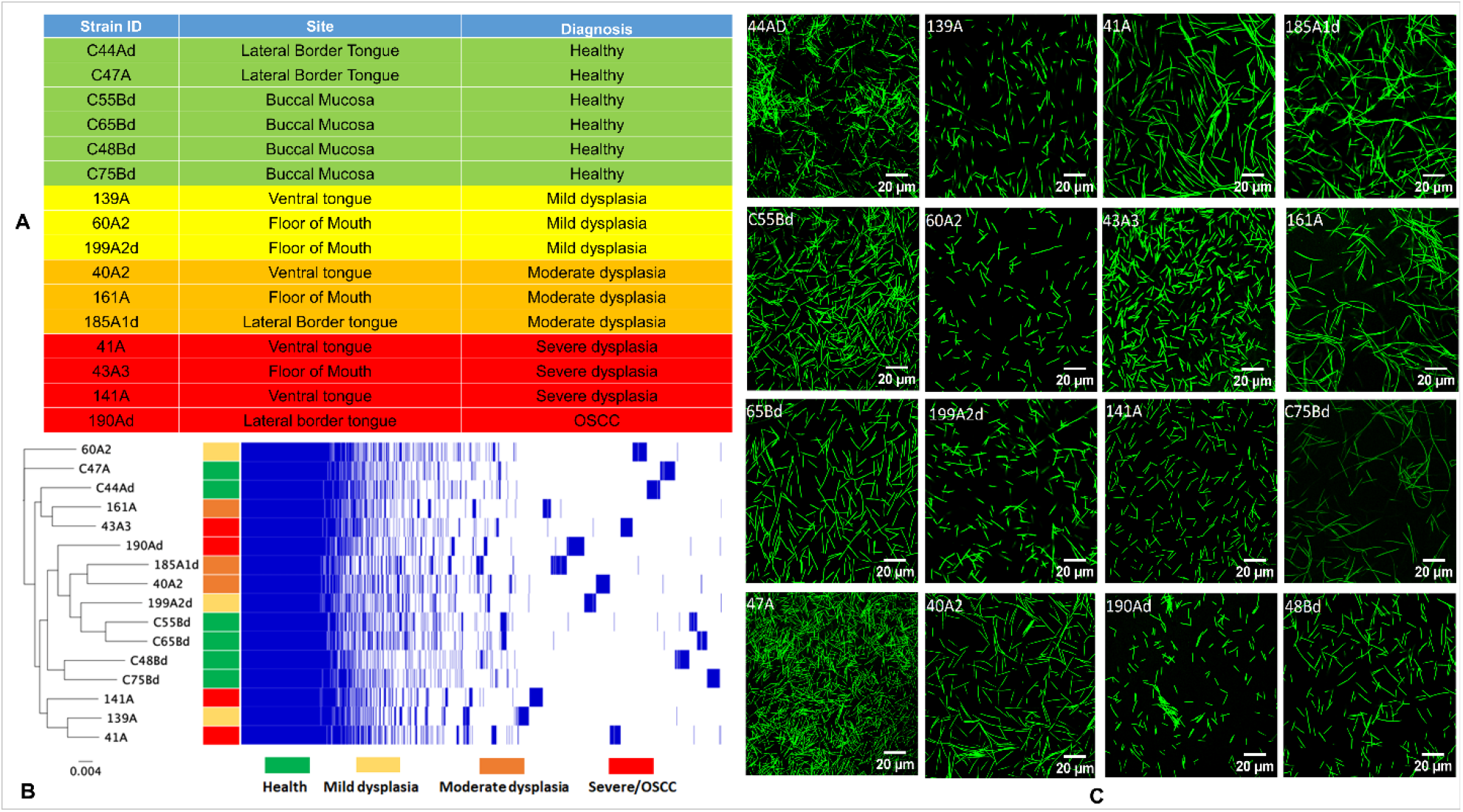
The clinical isolates of *F. polymorphum* included in the study. **A)** Sixteen representative *F. polymorphum* isolates isolated from the oral mucosa of healthy individuals or from the lesion surface of OLK patients with varying degrees of epithelial dysplasia. **B)** Phylogenetic tree generated from a core genome alignment with a matrix plot showing gene presence/absence. No distinct clustering of isolates by disease group was discerned based on core genomes (Similar results obtained with accessory genomes—see Supplementary Figure 1). **C)** Representative images of the 16 isolates by confocal microscopy—the bacteria were stained by SYTO 9 dye.

### Bacterial culture and inoculum preparation

For each experiment, the *F. polymorphum* isolates were revived from stock on brain heart infusion (BHI) agar plates supplemented with yeast extract, cysteine, and 5% sheep blood and then subcultured in BHI broth supplemented with yeast extract, cysteine and 1% vitamin K/hemin mix (Thermo Fisher Scientific, USA). Broth cultures were subsequently re-subcultured and grown to either exponential phase (OD_600_ ≈ 0.7–0.8) or stationary phase (OD_600_ ≈ 1.3–1.4), depending on the downstream assay. All cultures were maintained under strict anaerobic conditions (80% N_2_, 10% H_2_, 10% CO_2_). Contamination check on aerobic blood agar was performed regularly.

### Cell line authentication and culture

The dysplastic oral keratinocyte (DOK) cell line (37) was obtained from Sigma-Aldrich (USA) and supplied with a certificate of authentication. Further authentication was performed using in-house short tandem repeat (STR) analysis, and cultures were regularly tested and confirmed negative for mycoplasma contamination. DOK were maintained in Dulbecco’s Modified Eagle’s Medium (DMEM; Gibco) supplemented with 10% fetal bovine serum (FBS), 2.5 mM L-glutamine, 5 µg/mL hydrocortisone, and 1% penicillin-streptomycin. Cells were cultured at 37 °C in a humidified atmosphere of 5% CO2 and passaged at ∼80% confluence using standard trypsinization protocols. All experiments were performed within the first 10 passages.

### Proliferation assay

The effect of *F. polymorphum* isolates on cell proliferation was assessed using the Alamar Blue assay. Bacterial inocula were prepared by growing to exponential phase (OD_600_ ≈ 0.7–0.8). DOK were seeded into black 96-well plates at 3,000 cells/well in serum- and antibiotic-free medium and allowed to adhere overnight under serum-starved conditions. The medium was replaced with fresh medium containing serum but no antibiotics, and cells were infected with each isolate at a multiplicity of infection (MOI) of 10 for 4 h. After infection, wells were washed and replenished with medium containing serum and antibiotics. Uninfected wells and medium-only controls were included.

Proliferation was measured at 24, 48, and 72 h post-infection. At each time point, 20 μL of Alamar Blue reagent (Thermo Fisher Scientific) was added per well, plates were incubated for 4 h, and fluorescence was measured (excitation 530 nm, emission 590 nm, gain 30). Each assay was performed in two independent biological replicates with three technical replicates per strain.

### Migration and invasion assays

Cell migration and invasion were assessed using QCM™ Fluorimetric 96-well assays (ECM510 and ECM555, respectively, Sigma-Aldrich, USA) according to the manufacturer’s instructions with modifications. DOK were grown to ∼80% confluence, serum-starved overnight, and harvested into quenching medium. Cells were seeded into assay inserts at 5 × 10^4^ cells/well for migration or 1 × 10^5^ cells/well for invasion. Two experimental setups were used for each of the assays. In the first, DOK were infected directly in the inserts with the *F. polymorphum* isolates (grown to exponential phase) at an MOI of 10 for 4 h, after which infection was stopped by adding penicillin-streptomycin; medium containing 10% FBS served as the chemoattractant in the lower chamber. In the second setup, seeded DOK were left uninfected while supernatants from 24-h infected DOK cultures were placed in the lower chamber as chemoattractants. In both assays, plates were incubated for 16 h after infection.

Following incubation, migrated or invaded cells were detached, lysed, and stained according to the kit protocol. Fluorescence was measured using 480/520 nm filter set (gain = 65). Standard curves were generated from serially diluted DOK (20,000 to 625 cells/well) included in each plate. Each assay was performed in two independent experiments, with three technical replicates per isolate.

### Amyloid-like FadA Quantification

The clinical isolates of *F. polymorphum* were cultured for 36 hours to stationary phase (OD_600_ ≈ 1.3– 1.4). Cultures were then centrifuged at 3,500 × *g* for 5 minutes, washed twice in PBS, and resuspended to OD_600_ = 2. Amyloid-like FadA was measured using a Congo Red (CR) depletion assay as previously described (16). Briefly, 5 μL of CR stock solution (2 mg/mL) was added to 1 mL of bacterial suspension to achieve a final concentration of 10 μg/mL. After incubation at room temperature for 10 minutes, suspensions were centrifuged at maximum speed for 10 minutes, and 200 μL of each supernatant was transferred to a 96-well plate for OD500 measurement. Amyloid-like FadA levels were inferred from the reduction in supernatant OD relative to CR-only controls. Each isolate was tested in triplicate in three independent experiments.

### Transcriptomic analysis of *F. polymorphum* clinical isolates

The clinical isolates of *F. polymorphum* were cultured to mid-log phase (OD_600_ ≈ 0.7–0.8). To stabilize RNA, stop solution (95% absolute ethanol, 5% phenol, v/v) was added to cultures at a 1:5 ratio, mixed thoroughly, and immediately centrifuged (3,500 × *g*, 5 min, 4 °C). Pelleted cells were processed for total RNA extraction using the PureLink™ RNA Mini Kit (Thermo Fisher Scientific, USA) according to the bacterial RNA workflow, with the following modifications: homogenization was achieved by pipetting rather than mechanical disruption. Eluates were stabilized with SUPERase•In™ RNase inhibitor (Thermo Fisher Scientific, USA) followed by DNase treatment (TURBO DNA-free™ kit, Thermo Fisher Scientific, USA).

RNA integrity analysis, library construction (rRNA depletion) and Illumina sequencing (NovaSeq X Plus, 2 × 150 bp) and standard bioinformatic analysis were performed by Novogene (USA). Genome assembly ASM15362v1 from *F. polymorphum* ATCC 10953 was used as a reference. To investigate isolate-level transcriptional variation, each isolate was assigned a severity score based on the clinical category of its origin (1 = health, 2 = mild dysplasia, 3 = moderate dysplasia, 4 = severe dysplasia/OSCC). Spearman correlation analysis was then used to identify bacterial genes whose expression significantly correlated with severity score (|rs| ≥ 0.5, p ≤ 0.05).

### Transcriptome and cytokine profiling of DOK

#### Co-culture set-up

DOK were seeded in 24-well plates (1.5 × 10^5^ cells/well) and serum-starved overnight in antibiotic-free medium. On the following day, cells were infected with the clinical *F. polymorphum* isolates at an MOI of 10 for 4 h. Medium was then replaced with serum-containing medium, and cells were incubated for an additional 20 h. Uninfected cells served as controls. Conditioned media were collected, centrifuged, and filter-sterilized for cytokine analysis, while cell monolayers were processed for RNA extraction.

#### Cytokine analysis

Discovery multiplex cytokine quantification was performed at Eve Technologies (Calgary, Canada) using Luminex™ 200 system (Luminex, Austin, TX, USA). Ninety-six cytokines, chemokines, and growth factors were measured using the MILLIPLEX® MAP Human Cytokine Panels A and B (MilliporeSigma, USA). Two replicates per isolate were analyzed.

#### RNA extraction and sequencing

Total RNA was extracted using the PureLink™ RNA Mini Kit (Thermo Fisher Scientific, USA). Eluates were treated as described above for RNA from the isolates; RNA integrity was confirmed using a 2100 Bioanalyzer system (Agilent Technologies, USA). For each isolate, RNA from three technical replicates were pooled for analysis. Library preparation (poly-A enrichment), sequencing (NovaSeq X Plus, 2 × 150 bp), and primary bioinformatics analysis (quality control, mapping to reference genome, quantification, normalization, and differential expression analysis) were performed by Novogene (USA) according to standard protocols.

#### Differential gene expression and pathway enrichment analysis

RNA expression data from DOK cells were analyzed using DESeq2 (38). Pairwise differential expression analyses were performed to compare DOK responses among three conditions: uninfected controls, infection with health-derived isolates, and infection with disease-derived isolates. Genes with adjusted p-value (Benjamini– Hochberg) ≤ 0.05 and |log2 fold change| ≥ 1 were considered as significantly differentially expressed. They were then ranked by fold change and used as input for gene set enrichment analysis (GSEA) (39) against the Hallmark collection in the Molecular Signatures Database (MSigDB v7.1) setting false-discovery rate (FDR) at ≤ 0.1.

#### Correlation-based analysis

To capture associations with lesion severity, each experimental condition was assigned a severity score based on isolate origin (0 = no infection, 1 = health, 2 = mild dysplasia, 3 = moderate dysplasia, 4 = severe dysplasia/OSCC). Spearman correlation was used to identify genes significantly associated with severity (|rs| ≥ 0.5, *p* ≤ 0.05). Significantly correlating genes were then ranked by fold change and analyzed by GSEA (MSigDB v7.1), using an FDR cutoff of 0.2

### Validation of ANKRD26 expression

ANKRD26 was selected for validation as the top positively correlating gene with severity score. Gene expression was quantified using a predesigned TaqMan primer/probe set (Thermo Fisher Scientific, USA) in a one-step reaction as previously described (40), with GAPDH as the reference gene. The same RNA samples used for RNA-seq were analyzed. Western blot analysis was performed in DOK infected with four representative *F. polymorphum* isolates (one per clinical category) and in uninfected controls. DOK (3 × 10^5^/well, 12-well plates) were infected at MOI 10 overnight. Cells were lysed in RIPA buffer with protease inhibitors, and total protein was quantified (Qubit Protein Assay, Thermo Fisher Scientific). Equal protein amounts (30–40 µg) were separated on 7.5% SDS-PAGE gels and transferred to PVDF membranes. Blots were probed with anti-ANKRD26 clone A302-307A (Bethyl Laboratories, 1:1000) and anti-β-actin (Cell Signaling, 1:5000), followed by HRP-conjugated secondary antibodies. Detection was performed by enhanced chemiluminescence (ECL, Thermo Fisher Scientific) on a ChemiDoc system (Bio-Rad).

### Validation of infection-correlated host genes in TCGA OSCC dataset

To validate the relevance of DOK genes that correlated with infection severity score, transcriptomic data from The Cancer Genome Atlas (TCGA) Head and Neck Squamous Cell Carcinoma cohort were analyzed. Clinical data were used to restrict the dataset to OSCC subsites (n = 243 primary tumor samples, 44 solid tissue normal controls). RNA expression data were downloaded from the Genomic Data Commons (GDC) and processed using the DeSeq2 pipeline. Genes were considered significantly differentially expressed if they had an adjusted p-value (Benjamini–Hochberg) ≤ 0.05.

A total of 80 host genes that showed the strongest correlation with the clinical category of *F. polymorphum* isolates *in vitro* (top 40 positively and 40 negatively correlated at 4 and 24 hours post-infection of DOK) were selected for validation. A match was defined as a gene significantly differentially expressed in tumors versus normal tissues in the same direction as observed in the *in vitro* infection experiments.

## Results

We selected 16 isolates of *F. polymorphum* from a collection of isolates previously recovered from the oral cavities of patients with varying levels of dysplasia or healthy controls; one isolate from OSCC was pooled with those from severe dysplasia (**Fig. 1A**). Isolate relatedness was assessed following phylogenetic analysis of an alignment of the core genome, including 1,245 shared genes (**Fig. 1B**). No relationship between *F. polymorphum* genotype and disease severity was identified. A phylogeny based on the accessory *F. polymorphum* genome showed a similar lack of association (**Supplementary Fig. 1**). A genome-wide association analysis using SCOARY did not identify any *F. polymorphum* genes that were unique to the disease-derived isolates or severe-dysplasia/OSCC-derived isolates. Phenotypically, the isolates exhibited a typical tapered rod or fusiform morphology, although the length of the individual rods varied between isolates (**Fig. 1C**).

### Clinical isolates of *F. polymorphum* modulate DOK phenotypes in a dysplasia severity– dependent manner

To determine whether the source of *F. polymorphum* isolates influences their interaction with host cells, we evaluated the effects of clinical isolates from health and disease on dysplastic oral keratinocytes (DOK). For proliferation, DOK were infected with each isolate at MOI 10 and proliferation was assessed at 24, 48, and 72 h post-infection using the Alamar Blue assay. Health-derived isolates had no significant effect on cell growth, whereas disease-derived isolates induced a marked increase (**Fig. 2A**). Strikingly, isolates from patients with mild, moderate, and severe dysplasia/OSCC—hereafter “high-risk” isolates—elicited a stepwise increase in DOK proliferation, with strong correlations to disease severity observed at 48 and 72 h (**Fig. 2B–C**).

**Figure 2.**
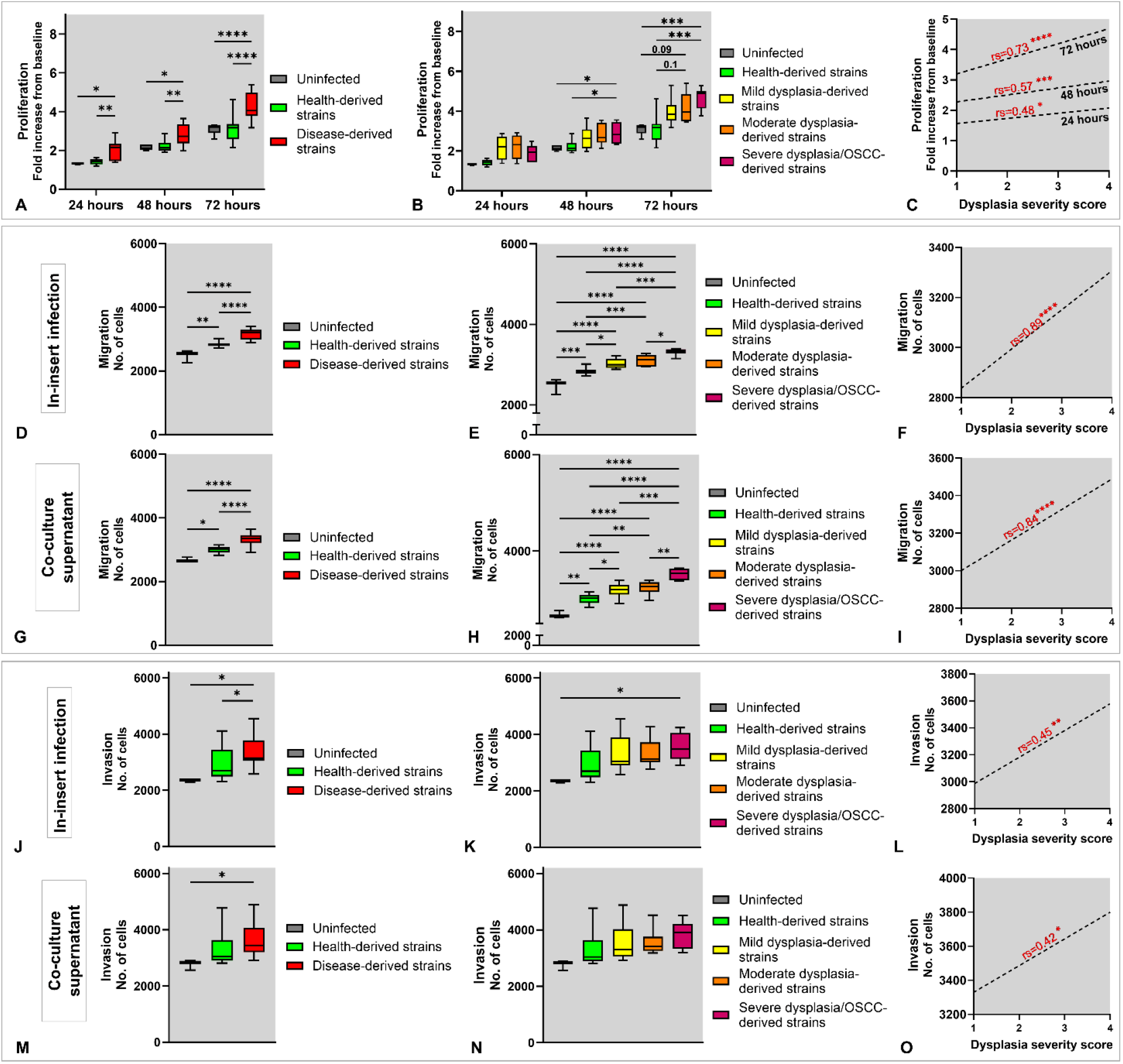
Proliferation, migration and invasion of DOK in response to infection with clinical isolates of *F. polymorphum*. For proliferation (upper panels), dysplastic oral keratinocytes (DOK) were infected with each isolate at an MOI of 10 or left uninfected for four hours, and proliferation was assessed at 24, 48, and 72 hours post-infection using the Alamar Blue assay. **A)** Disease-derived isolates vs. health-derived isolates. **B)** Pairwise comparisons based on clinical categories of the isolates. **C)** Correlation with dysplasia severity score (1 to 4 for healthy, mild, moderate and severe dysplasia/OSCC, respectively). Transwell migration assays (middle panel) were performed using two experimental setups: **D-F)** DOK were infected directly in the inserts while culture medium containing 10% FBS was used as attractant; and **G-I)** Uninfected DOK were exposed to supernatants from 24-hour co-cultures as attractants. Panels D–F and G–I reflect comparisons/correlations as in A–C. Transwell invasion assays were performed similarly: **J–L)** In-insert infection and **M–O)** Supernatant-attractant. Each plot represents data from two independent experiments, each in three technical replicates for each isolate. Statistical significance was determined using two-way ANOVA with post hoc Tukey’s test for group comparisons and Spearman correlation (rs) for correlations, and significance is indicated as follows: p ≤ 0.05 (*), p ≤ 0.01 (**), p ≤ 0.001 (***), and p ≤ 0.0001 (****).

Transwell migration assays showed that either direct *F. polymorphum* infection (**Fig. 2D**) or exposure to supernatants of *F. polymorphum-*DOK co-cultures (**Fig. 2G**) significantly promoted migration of DOK but the effect was significantly greater with high-risk isolates. Stratification by dysplasia grade revealed that isolates from more severe lesions induced progressively stronger migration responses in both experimental setups (**Fig. 2E–I**). Transwell invasion assays were performed using ECMatrix™-coated inserts under the same two experimental setups as for migration. High-risk isolates significantly enhanced invasion, whereas health-derived “low-risk” isolates had little or no effect (**Fig. 2J, M**). As with proliferation and migration, invasion responses correlated with the dysplasia severity of the isolate’s patient of origin, with isolates from higher-grade lesions eliciting the strongest invasive responses (**Fig. 2K–O**).

### Amyloid-like FadA production correlates with isolate-specific effects on DOK cells

Although low- and high-risk *F. polymorphum* isolates were genotypically similar, they elicited strikingly different responses in DOK cells. To determine whether phenotypic variation might account for these differences, we examined FadA, a key amyloid-like virulence factor implicated in the carcinogenic activity of *Fusobacterium*. Quantitative analysis using a Congo Red depletion assay showed that high-risk isolates exhibited significantly greater Congo Red absorption (reflecting higher FadA expression) compared to low-risk isolates (**Fig. 3A**). Importantly, when isolates were stratified by the dysplasia severity of the patient of origin, Congo Red binding increased stepwise, with the lowest levels in health-derived isolates and the highest in severe dysplasia/OSCC, demonstrating a strong correlation with severity score (**Fig. 3B–C**).

**Figure 3.**
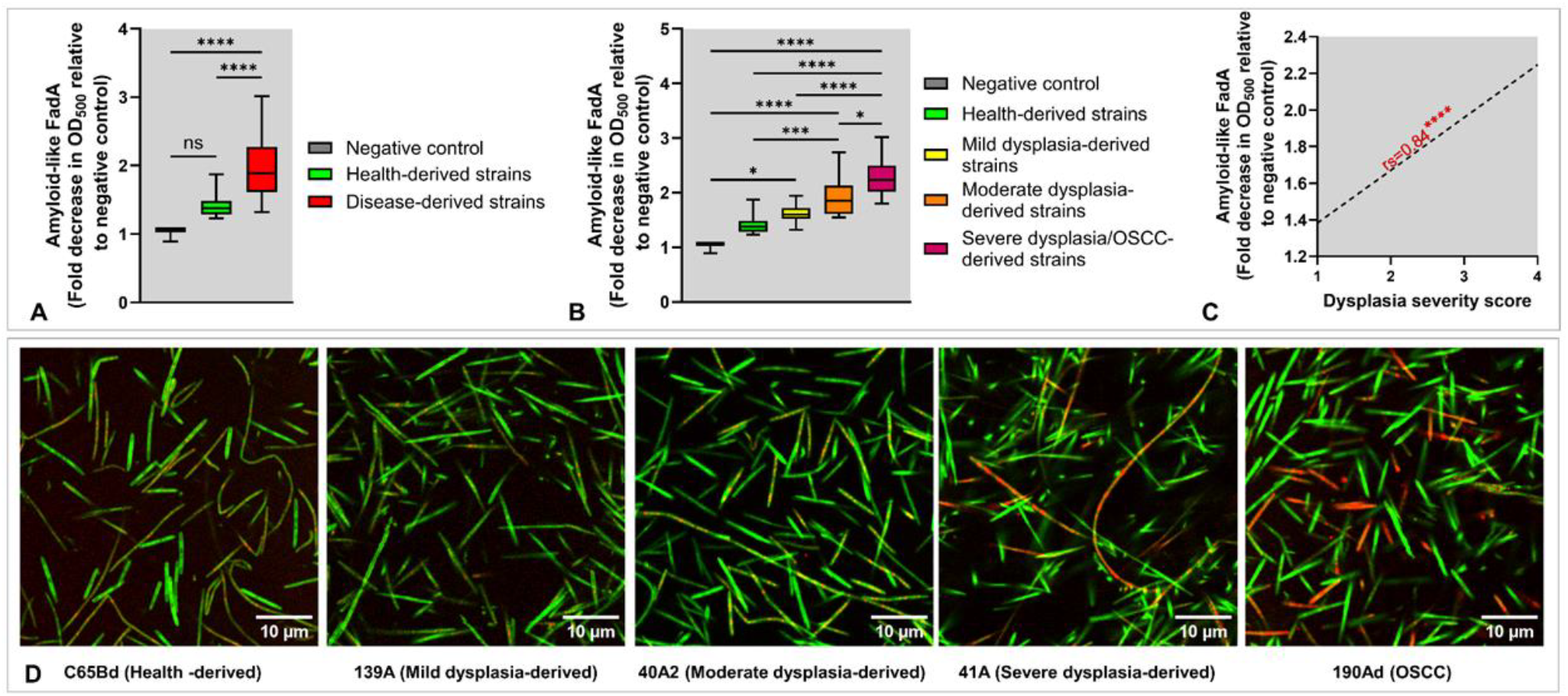
Quantification of amyloid-like FadA in clinical isolates of *F. polymorphum*. Sixteen isolates were grown to stationary phase (36 hours). Amyloid-like FadA was quantified using a Congo Red (CR) depletion assay as described in Methods). **A)** Disease-derived isolates vs. health-derived isolates. **B)** Pairwise comparisons based on clinical categories of the isolates. **C)** Correlation with dysplasia severity score (1 to 4 for healthy, mild, moderate and severe dysplasia/OSCC, respectively). **D)** Confocal microscopy of FadA (CR) and representative bacterial isolates (SYTO 9). Each plot represents data from three independent experiments, each in three technical replicates for each isolate. Statistical significance was determined using one-way ANOVA with post hoc Tukey’s test for group comparisons and Spearman correlation (rs) for correlations, and significance is indicated as follows: p ≤ 0.05 (*), p ≤ 0.01 (**), p ≤ 0.001 (***), and p ≤ 0.0001 (****).

### Expression of heme uptake genes in *F. nucleatum* isolates correlates with dysplasia severity

To explore whether additional bacterial virulence factors might contribute to isolate-specific differences, we analyzed the transcriptomes of the *F. polymorphum* isolates for genes associated with the dysplasia severity of the host. Spearman correlation analysis (severity score 0–4) identified multiple genes significantly correlated with increasing severity, with three of the top five correlating genes involved in heme uptake (**Fig. 4A**). Heatmap analysis further confirmed that these heme acquisition genes were most highly expressed in high risk isolates, particularly those from moderate and severe dysplasia/OSCC cases (**Fig. 4B**).

**Figure 4.**
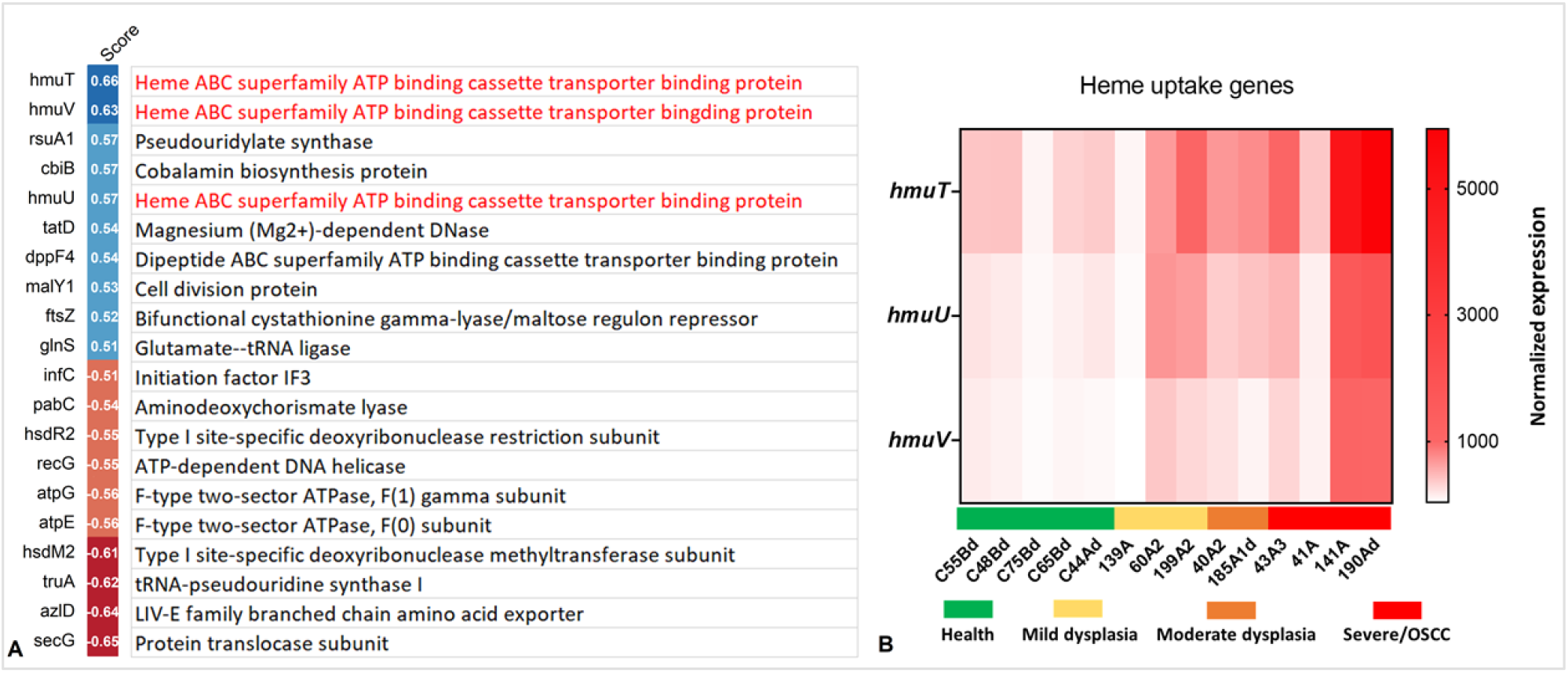
Transcriptional profiling of *F. polymorphum* clinical isolates. Sixteen isolates were grown to mid-log phase (OD500 ∼ 0.7) followed by total RNA extraction for mRNA sequencing. Isolates were assigned a severity score from 0 to 4 based on their origin, 1 = health, 2 = mild dysplasia, 3 = moderate dysplasia, and 4 = severe dysplasia/OSCC. Spearman correlation analysis was performed to identify genes significantly correlating with the severity score (|rs| ≥ 0.5, p ≤ 0.05). **A)** Top positively and negatively correlated genes. **B)** Expression heatmap of three heme uptake genes (*hmuT, hmuV hmuU*) across isolates grouped by clinical origin.

### Variation in DOK responses is primarily explained by proliferative signaling on a background of consistent inflammatory and metabolic responses

To investigate the molecular responses underlying variable responses of DOK to the *F. polymorphum* isolates, we performed mRNA-seq and multiplex cytokine analysis following infection with the isolates at MOI 10:1 for 24 hours. Standard differential gene expression analysis revealed highly similar of DOK across *F. polymorphum* isolates, regardless of clinical category. Specifically, both low- and high-risk isolates induced robust enrichment of NF-κB, “inflammatory response,” and the inflammatory arm of KRAS signaling Hallmark gene sets, while consistently downregulating oxidative phosphorylation (**Fig. 5A–B**). These transcriptomic signatures were consistent with cytokine profiling of culture supernatant, which revealed a shared secretion pattern across isolates dominated by IL-8, CXCL1, CXCL16 and IL-6 (**Fig. 5C**). Comparable findings were also observed after 4 hours of infection (**Supplementary Fig. 2**).

**Figure 5.**
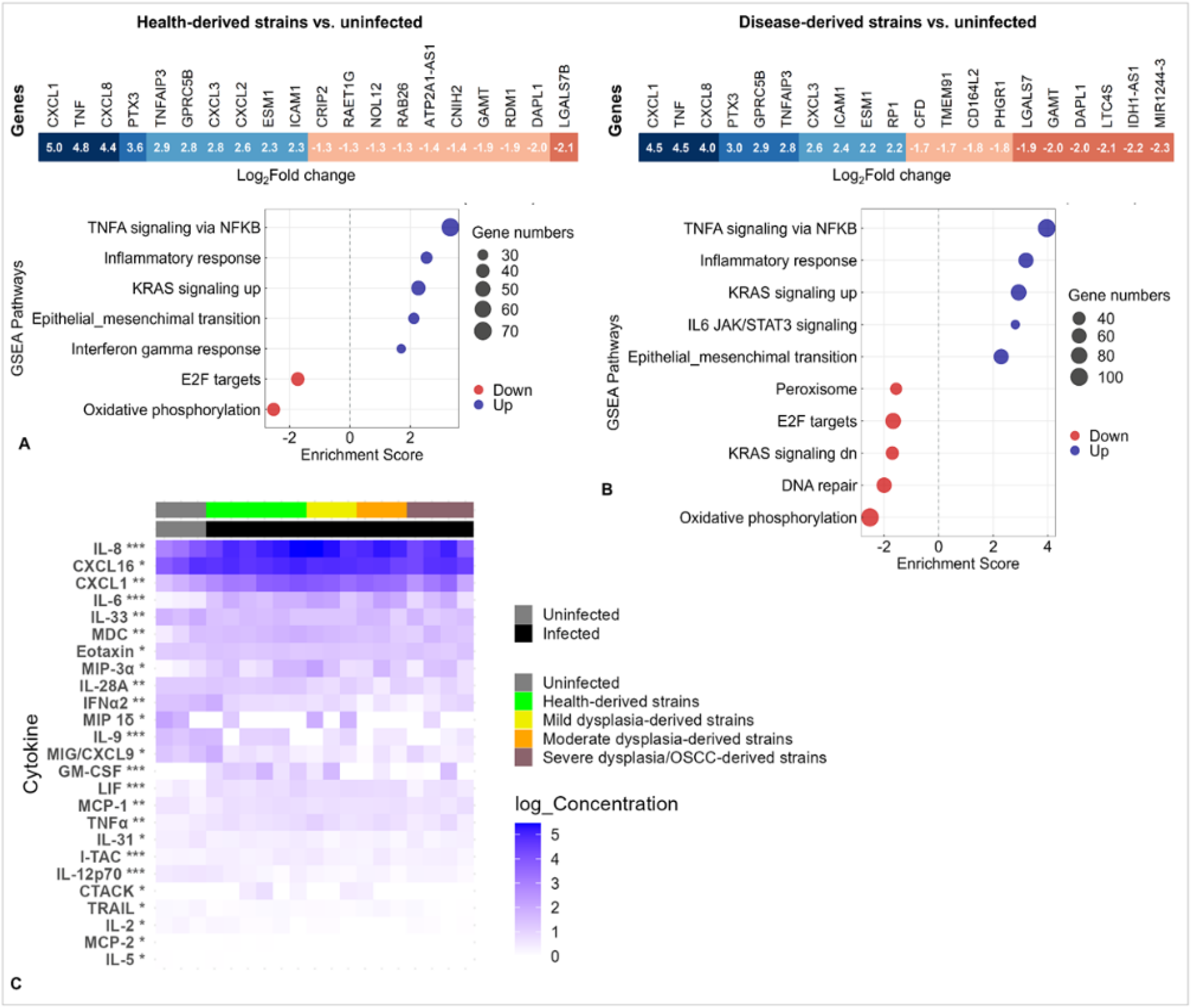
Transcriptional and cytokine responses of DOK to infection with clinical isolates of *F. polymorphum*. DOK were infected at an MOI of 10 or left uninfected for four hours followed by 20-hour incubation. RNA was extracted from cell monolayers for mRNA-seq (pooled RNA from 3 technical replicates/isolate). Co-culture supernatants were collected for discovery multiplex cytokine analysis (two technical replicates per isolate). **A)** Top up- and downregulated genes (FDR ≤ 0.1) and significantly enriched pathways (FDR ≤ 0.1) in DOK infected with health-derived isolates. **B)** Same as (A) but for disease-derived isolates. **C)** Heatmap of cytokines and chemokines significantly increased across all isolates independent of severity (infection vs. no infection) (t-test; * FDR ≤ 0.25, ** FDR ≤ 0.1, *** FDR ≤ 0.05).

In contrast, when we examined transcriptional responses in relation to the dysplasia severity of the patient of origin (severity scores 0–4), we found clear evidence of variation. Spearman correlation analysis identified genes significantly associated with increasing severity, with the top correlations presented in **Fig. 6A**. Pathway enrichment analysis of these severity-associated genes identified predominantly upregulation of proliferative signaling, including enrichment of mitotic spindle and G2M checkpoint gene sets, besides further downregulation of oxidative phosphorylation and DNA repair (**Fig. 6B**). Although NF-κB/TNFα and “inflammatory response” pathways were also enriched, the leading-edge genes (REL, TRIB1, MYC, ETS2) reflected regulatory and proliferative factors rather than canonical cytokines, which were uniformly induced across all isolates.

**Figure 6.**
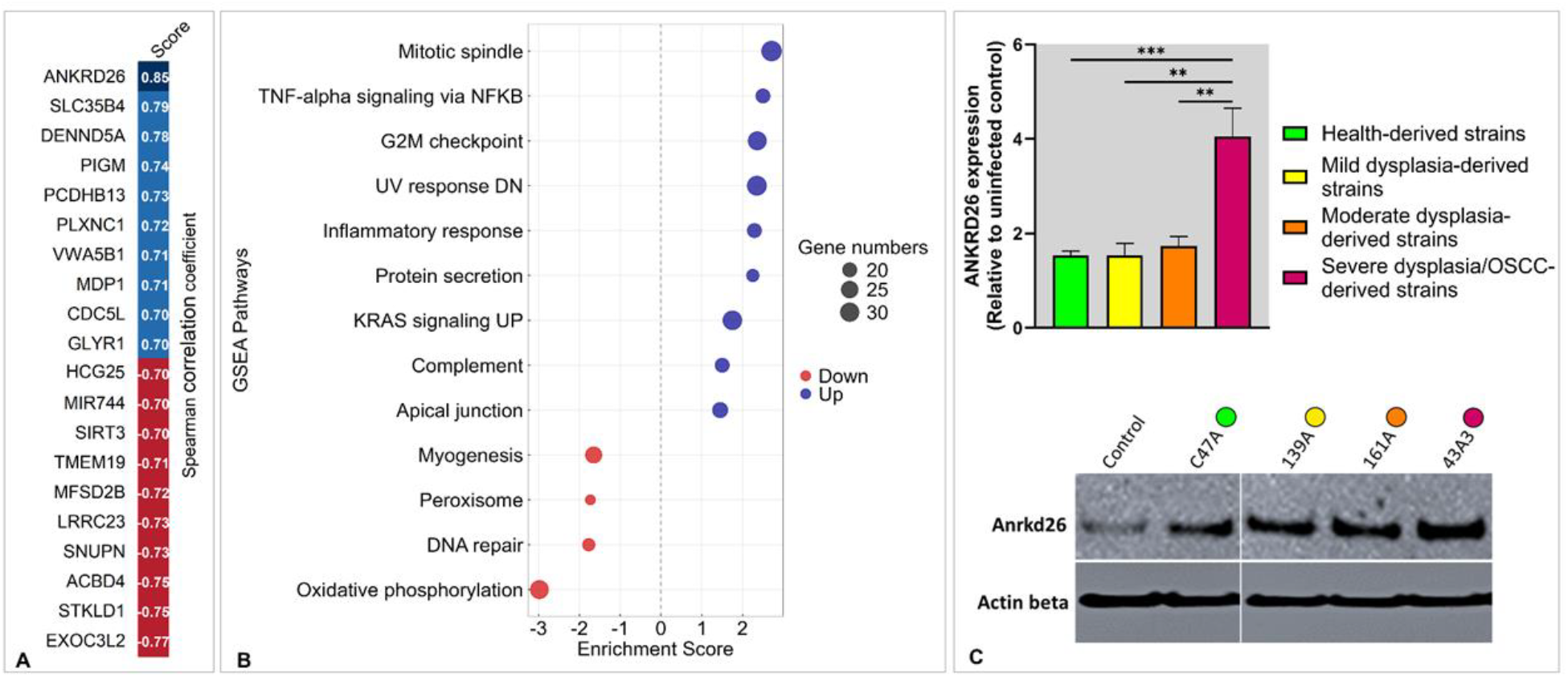
Correlation-based analysis of transcriptional responses to *F. polymorphum* infection. DOK were infected or left uninfected. Total RNA was extracted at 24 hours post-infection for mRNA sequencing (pooled RNA from 3 technical replicates/isolate). Each condition was assigned a severity score based on clinical category of the isolate (0 = no infection, 1 = health, 2 = mild dysplasia, 3 = moderate dysplasia, and 4 = severe dysplasia/OSCC. Spearman correlation analysis was performed to identify genes significantly correlating with the severity score (|rs| ≥ 0.5, p ≤ 0.05). **A)** Top positively and negatively correlated genes. **B)** Pathways significantly enriched (FDR ≤ 0.2) in correlation with increasing severity score. C) Validation of *ANKRD26* by qPCR (same RNA samples) and Western blot (independent experiment). Statistical significance (qPCR only) was determined using one-way ANOVA with post hoc Tukey’s test for group comparisons: p ≤ 0.01 (**), and p ≤ 0.001 (***).

To validate severity-linked responses, expression of the top correlating gene, *ANKRD26*, was confirmed by qRT-PCR and Western blot analysis, both showing increased expression in DOK infected with severe dysplasia/OSCC-derived isolates (**Fig. 6C**).

### Genes correlating with infection by clinical *F. polymorphum* isolates mirror expression trends in TCGA OSCC dataset

To assess the clinical relevance of our in vitro results, we compared the *F. polymorphum* infection– correlated gene set with expression profiles from the TCGA OSCC dataset. Specifically, we examined the 80 genes most strongly correlated with isolate severity (the top 40 positively and 40 negatively correlated at 4 and 24 hours post-infection of DOK; **Fig. 7**). Of these, 36 genes (45%) were significantly differentially expressed in OSCC tumors compared with normal tissues (*padj* < 0.05) in the **same direction** as in our in vitro assays—that is, genes positively correlated with dysplasia severity in infection experiments were upregulated in tumors, while negatively correlated genes were downregulated.

**Figure 7.**
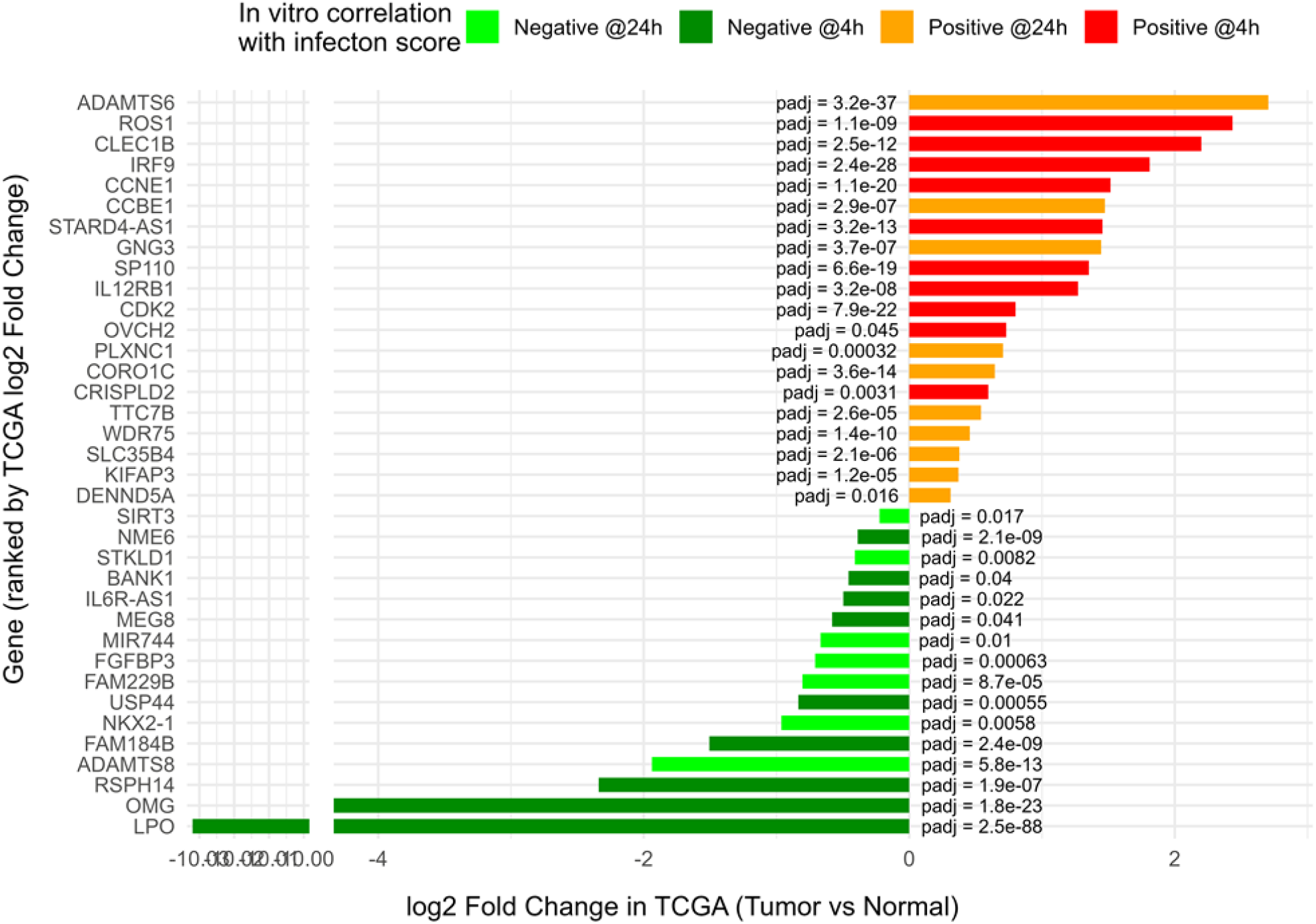
Validation of *F. polymorphum* infection-correlated genes in the TCGA OSCC dataset. Eighty genes showing the strongest correlation with *F. polymorphum* clinical category (top 40 positively and 40 negatively correlated at 4 and 24 hours post-infection of DOK), as described in Figure 6, were examined in TCGA oral squamous cell carcinoma (OSCC) samples (n = 243 primary tumors, 44 solid tissue normal controls). Thirty-six genes (45%) were significantly differentially expressed (padj < 0.05) in tumors compared to normal tissues in the same direction as in the in vitro infection. Bar plot shows TCGA log2 fold changes for each gene, colored by *in vitro* correlation category (positive or negative at 4 h or 24 h). Adjusted p-values are shown beside the bars. Differential expression analysis in TCGA was performed using DESeq2.

## Discussion

Most evidence on the carcinogenicity of *F. nucleatum* comes from studies on CRC (41); its role in OSCC remains understudied. Our previous microbiome analyses identified *F. polymorphum* as the most prevalent *Fusobacterium* lineage associated with oral leukoplakia (OLK) (31). We also showed that *F. polymorphum*-linked OTUs are enriched in OLKs with severe dysplasia and in OSCC (1, 20). Yet, *F. polymorphum* is also a common commensal of healthy mucosa (25, 26), and pangenome analysis revealed no genotype specifically linked to dysplasia (31). This raised the question of whether strain-level phenotypic heterogeneity might account for its role in malignant progression.

Our current findings support this hypothesis. Isolates from dysplastic lesions consistently enhanced DOK proliferation, migration and invasion, and the magnitude of these effects correlated with the severity of dysplasia from which the isolates were obtained. In contrast, health-derived isolates showed weaker effects. This is the first demonstration of isolate-specific oncogenic differences within oral *Fusobacterium* species, establishing the novel concept of “high-risk” versus “low-risk” isolates in oral carcinogenesis. Importantly, genomic analyses did not reveal isolate-specific genotypes or accessory genes associated with carcinogenicity. Instead, phenotypic variation was closely tied to amyloid-like FadA expression.

FadA is a well-established virulence factor of *F. nucleatum*, known to form amyloid like aggregates that bind E-cadherin and activate β-catenin signaling via Annexin A1, thereby promoting carcinogenesis (12, 16). Our study demonstrates a clear association between amyloid-like FadA levels and the differences observed in DOK responses to the isolates, supporting a model in which elevated FadA expression contributes to the pro-carcinogenic activity of high risk isolates. Notably, this phenotype was stable *in vitro* despite storage and repeated culturing, suggesting heritability rather than transient induction. Since promoter and gene sequencing did not reveal differences (**Supplementary Fig. 3**), we hypothesize that stress-induced epigenetic regulation could drive increased translation or secretion of FadA in high risk isolates. The concomitant upregulation of heme acquisition genes in these isolates may reflect a stress-adapted phenotype, potentially linked to the inflamed and iron-limited microenvironment of dysplastic epithelium. Metatranscriptomic analysis of OSCC also confirms that *F. nucleatum* exhibits increased expression of iron acquisition genes in tumors *in vivo* (21). These findings provide further evidence for presence of genome-independent phenotypic features that shape carcinogenicity of *F. polymorphum*.

At the host side, cytokine and transcriptomic responses of DOK were broadly similar between low- and high-risk isolates, characterized by NF-κB activation, inflammatory mediators, and EMT-related genes. However, correlation-based analyses revealed that proliferation-associated pathways (mitotic spindle, G2M checkpoint) were specifically enriched in DOK infected with high-risk isolates. These data suggest that while inflammatory and metabolic responses are conserved across isolates, proliferative signaling scale with dysplasia severity and may underlie isolate-specific phenotypic effects on DOK. This, however, contradicts with previous studies on CRC showing the increased migration and invasion are mediated by cytokines and chemokines, including IL-8 and CXCL1 (9). Interestingly, our experiments using conditioned medium as attractant reproduced differences in migration and invasion induced by infection, despite lack of differences in cytokines concentrations among the groups, suggesting that other soluble factors, either secreted by *F. polymorphum* itself or induced in DOK cells, may mediate these phenotypes. Indeed, a previous study showed that *F. nucleatum* mediates its carcinogenic properties through their outer membrane vesicles, which also contains FadA (42).

Among the genes most strongly correlated with high-risk *F. polymorphum*, ANKRD26 stood out as highly upregulated and was validated by qPCR and Western blot. ANKRD26 encodes a protein implicated in hematopoietic proliferation, and sustained expression prevents internalization of cytokine receptors, leading to cytokine hypersensitivity (43). Its upregulation in DOK infected with disease-associated isolates may contribute to both enhanced proliferation and cytokine responsiveness, which may, at least in part, explain the variation in DOK responses to the isolates.

Importantly, nearly half (45%) of the top infection-correlated genes identified *in vitro* were also differentially expressed in the same direction in tumors compared with normal tissue in the TCGA OSCC dataset. This overlap strongly supports the biological relevance of our *in vitro* findings and suggests that FadA-driven effects and associated transcriptional programs may also be active *in vivo*.

In conclusion, this study presents a new model whereby *F. polymorphum* colonization of dysplastic lesions promotes malignant phenotypes in keratinocytes, with disease-associated isolates exhibiting greater oncogenicity than health-derived strains, establishing, for the first time, the novel concept of “high-risk” versus “low-risk” isolates in oral carcinogenesis. This appears to be mediated genome-independent regulation of key virulence factors. We propose a “vicious cycle” model in which colonization of dysplastic epithelium by *F. polymorphum* leads to inflammation and stress, in turn inducing FadA expression, and possibly other virulence factors, which further drives dysplasia progression and tumorigenesis. Further studies *in vivo* validation experiments and multiomics epigenetic and phenotypic characterization of *F. polymorphum* are warranted.

## Acknowledgement

We thank Dr. Bettina Buttaro, Lewis Katz School of Medicine, Temple University, for assistance with confocal laser microscopy imaging.

