## Supplementary Fig. for "*In vitro* analysis of *Fusobacterium Polymorphum* from oral leukoplakia patients identifies “high-risk” isolates"

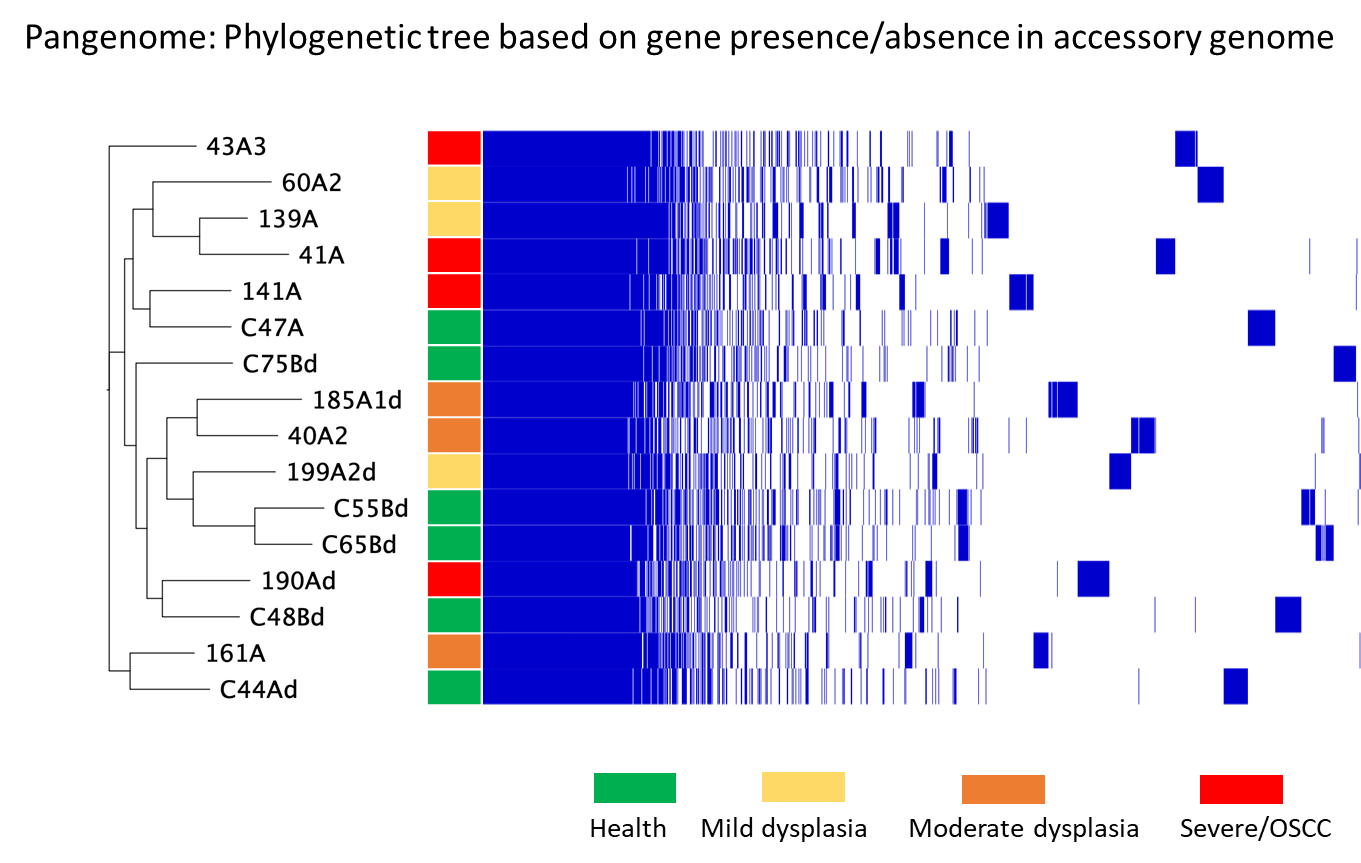

**Supplementary Figure 1. Accessory pangenome analysis.** Phylogenetic tree based on gene presence/absence in accessory genome

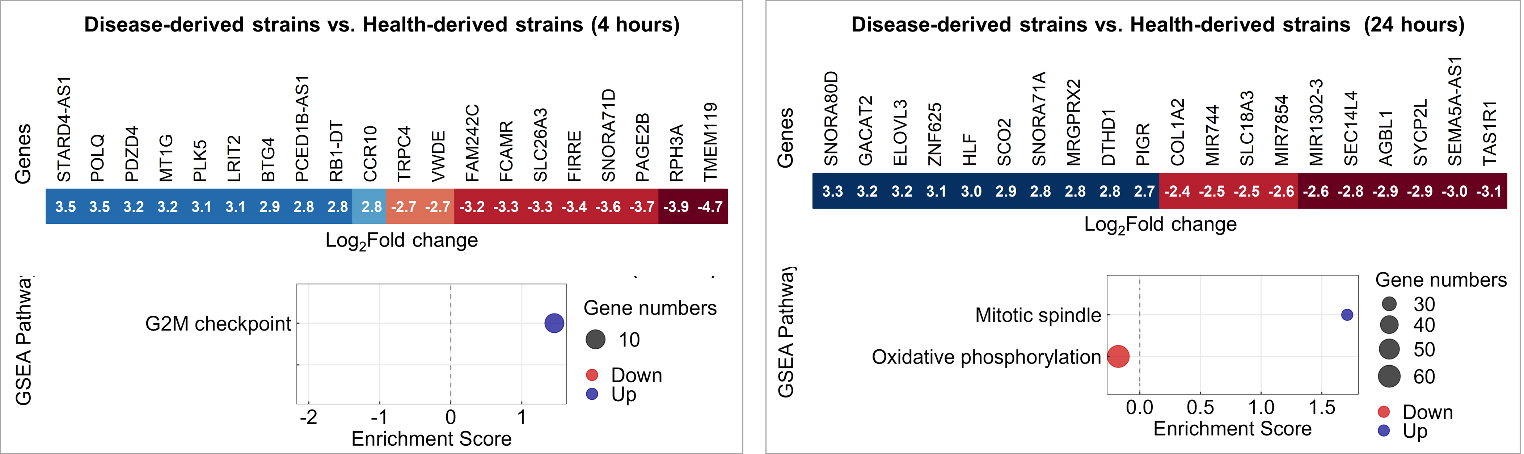

**Supplementary Figure 2. Transcriptional responses of DOK to infection with clinical isolates of *F. polymorphum* at 4 hours.**

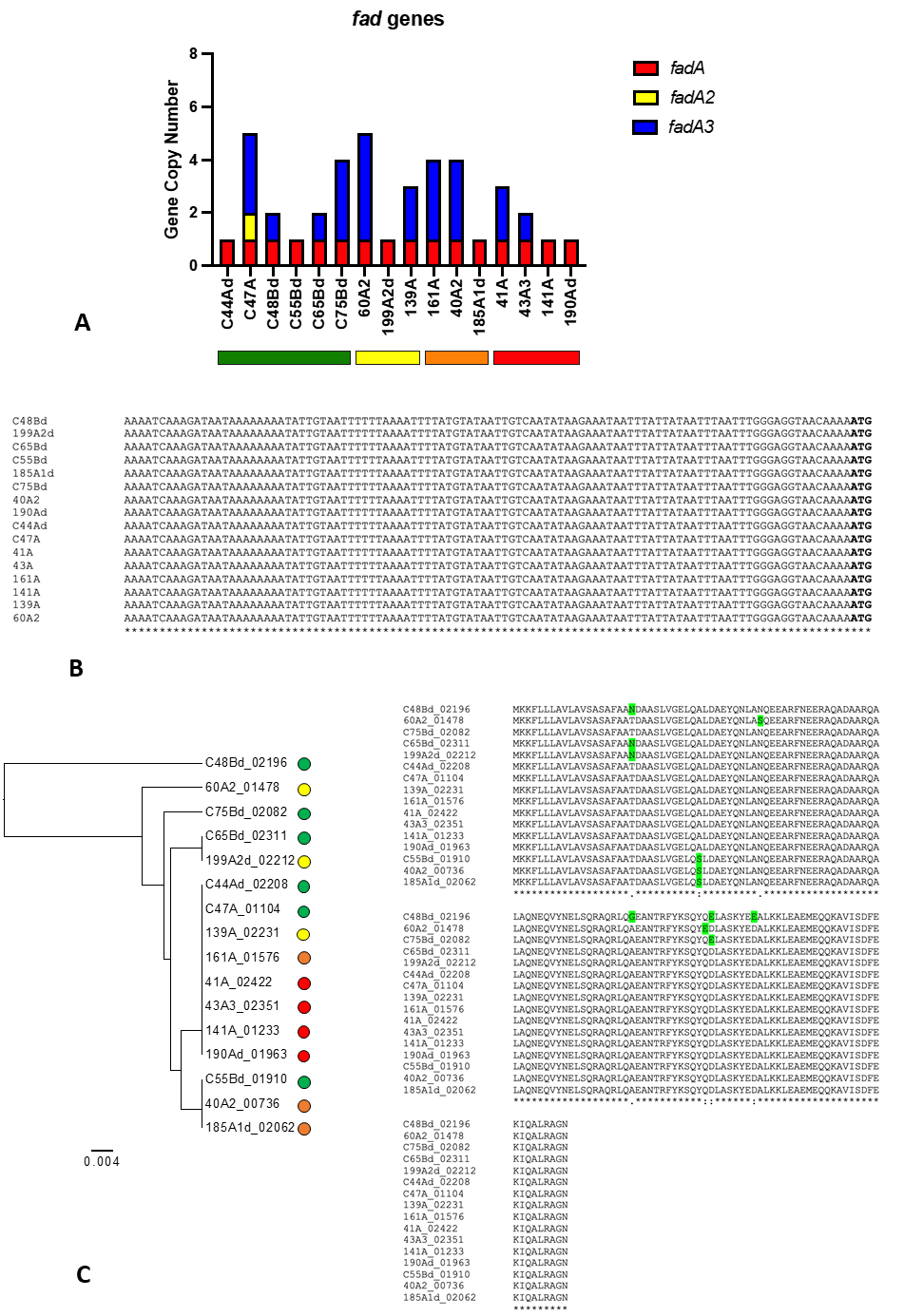

**Supplementary Figure 2. FadA gene analysis of 16 F. polymorphum isolates. A)** Gene copy number, B) promotor sequences, and C) gene sequences.
